# Dysregulated airway epithelial antiviral immunity in Down Syndrome impairs type III IFN response and amplifies airway inflammation during RSV infection

**DOI:** 10.1101/2024.11.22.624921

**Authors:** Elizabeth Chorvinsky, Surajit Bhattacharya, Betelehem Solomon Bera, Allison Welham, Karim Ismat, Claire M Lawlor, Diego Preciado, Jose L Gomez, Hiroki Morizono, Dinesh K Pillai, Maria J. Gutierrez, Jyoti K Jaiswal, Gustavo Nino

**Affiliations:** Division of Pediatric Pulmonary and Sleep Medicine, Children’s National Hospital, Washington, DC, USA; Center for Genetic Medicine Research, Children’s National Research Institute, Washington DC, USA; George Washington University School of Medicine and Health Sciences, Washington, D.C, USA; Division of Pediatric Otorhinolaryngology, Children’s National Hospital, Washington, DC; Pulmonary, Critical Care and Sleep Medicine Section, Yale University, New Haven, CT, USA; Division of Pediatric Allergy, Immunology and Rheumatology, Johns Hopkins University, Baltimore, MD, USA

**Keywords:** Down Syndrome, Trisomy 21, airway epithelial cell, RSV, bronchiolitis, interferon

## Abstract

Trisomy 21 (TS21), also known as Down syndrome (DS), increases pediatric mortality risk from respiratory syncytial virus (RSV) by nine-fold, yet its underlying immunological basis remains unclear. Here, we investigated RSV-induced immunological responses in TS21 airway epithelial cells (AECs), the primary site of respiratory virus entry and host defense. TS21 AECs exhibit hyperactive interferon (IFN) signaling and reduced RSV infectivity, but they also show impaired type-III IFN responses during viral infection. Furthermore, TS21 AECs demonstrate heightened production of proinflammatory mediators CXCL5 and CXCL10 both before and after RSV exposure. Infants with DS suffering from severe viral bronchiolitis demonstrate dysregulated airway immune responses in vivo, characterized by diminished type-III IFN levels and increased CXCL5/CXCL10 secretion. Our results indicate that RSV severity in DS is not due to impaired viral control but to dysregulated airway proinflammatory responses, offering new therapeutic opportunities to mitigate the severity of RSV infection in children with DS.

## INTRODUCTION

Down syndrome (DS) is a genetic condition resulting from trisomy of chromosome 21 (TS21) and is the most common chromosomal abnormality in humans^1^. Among individuals with DS, respiratory disorders pose a significant burden, leading to high morbidity and mortality across all age groups^2–5^. Young children with DS are severely impacted by respiratory infections^2–7^, particularly those caused by respiratory syncytial virus (RSV)^8,9^. A meta-analysis of over a million children that examined the risk of severe RSV disease in children with DS identified these individuals have approximately 9 times higher risk of RSV-related hospitalization and death^8^. Furthermore, relative to euploid children, patients with DS had worse respiratory outcomes including hospital stay duration, oxygen requirement, ICU admission, and the need for mechanical ventilation^8^. Thus, there is strong clinical evidence of an unexplained and impactful vulnerability to severe RSV infections among individuals with DS.

Despite the known clinical impact of DS on the severity of RSV infection, the mechanism by which TS21 causes this impact remains unclear. Notably, TS21 is linked to hyperactivation of interferon (IFN) signaling^10–12^, which is associated with the triplication of the four IFN receptor (IFNR) genes encoded on human chromosome 21 (Hsa21)^10–12^. This results in baseline JAK/STAT signaling hyperactivation and overexpression of IFN-stimulated antiviral genes^13,14^ that could protect against viral infection^15^. However, increased IFN signaling in TS21 cells is also reported to trigger excessive inhibitory feedback during viral exposure, reducing antiviral responses^11^. Thus, it remains unclear whether baseline IFN hyperactivation in DS would enhance or disrupt RSV infectivity in TS21 cells.

In addition to experiencing dysregulated IFN antiviral responses, individuals with DS also suffer from a global state of immune dysregulation involving multiple cellular components^16–20^. Recent studies have identified that people with DS exhibit a systemic cytokinopathy characterized by elevated levels of several cytokines and chemokines, collectively inducing a chronic proinflammatory state^18^. These abnormalities increase the risk of autoimmunity in individuals with DS^16–20^, and elevate their susceptibility to organ damage during infections due to overactive immune responses^16–20^.

While hyperactivation of IFN and pro-inflammatory responses in DS are documented in systemic immune cells^16–20^, their status in the airway epithelial cells (AECs) is lacking. This represents a critical gap since AECs serve as the point of entry and primary defense against respiratory viruses like RSV ^21^. To understand the biological implications of immune dysregulation in DS within these cells, we conducted comprehensive analyses of RSV-induced antiviral responses using primary AECs obtained from euploid and triploid pediatric donors. We have used single-cell transcriptomics, molecular profiling of JAK/STAT signaling activation, and assessment of antiviral and pro-inflammatory responses in vitro and in vivo in children with and without DS. Our studies revealed that, although TS21 AECs exhibit hyperactive IFN signaling and low RSV infectivity, they also suffer from reduced type-III IFN production in response to RSV infection. Moreover, both, prior to, and during RSV infection, TS21 AECs exhibit an exaggerated production of pro-inflammatory mediators linked to leukocyte/granulocyte chemotaxis (e.g., CXCL5 and CXCL10)^22,23^. These observations were validated in vivo in infants with DS that were suffering from severe viral bronchiolitis, where the dysregulated airway antiviral response was characterized by reduced type-III IFN levels and heightened secretion of proinflammatory chemokines. Collectively, these findings identify, for the first time, that the basis for the severe RSV infections in children with DS is not impaired control of viral infection in AECs. Instead, this is due to dysregulated AEC immune response, which favors proinflammatory changes over the generation of protective type-III IFN antiviral responses^24–26^. This discovery opens new therapeutic pathways focusing on the modulation of airway immunity to mitigate the severe impact of RSV infections in children with DS.

## RESULTS

### Airway epithelial cells of children with DS exhibit reduced level of RSV infection

There is strong evidence that children with TS21 have clinically more severe RSV disease ^8^. However, it remains uncertain whether this is primarily attributed to heightened RSV infectivity or the immune dysregulation of DS, which is increasingly recognized for its role in influencing various phenotypical features of this condition^16–20^. To address this fundamental question, we assessed the ability of RSV to infect nasal AECs from euploid and DS donors (n=12, **Fig. 1A, Supplementary Table 1**). Whole genome arrays confirmed the TS21 karyotype in all AEC cultures used (**Fig. 1B**). Using fluorescence microscopy to examine AECs infected with RSV encoding the GFP gene (RSV-GFP), we identified that TS21 donors had a decreased proportion of infected (GFP-expressing) AECs compared to euploid donors (**Fig. 1C**). To confirm these results, we used two complementary approaches - flow cytometry analysis to quantify the GFP expressing cells (**Fig. 1D-E**) and immunoblotting to quantify RSV genome encoded protein F levels in cell lysates (**Fig. 1F-G**). Examining time (5-24 hours) and dose responses using a range of RSV multiplicity of infection (MOI) (0.1-3) confirmed that TS21 AECs show reduced levels of RSV infection relative to euploid donors (**Fig. 1D-G**).

**Figure 1.**
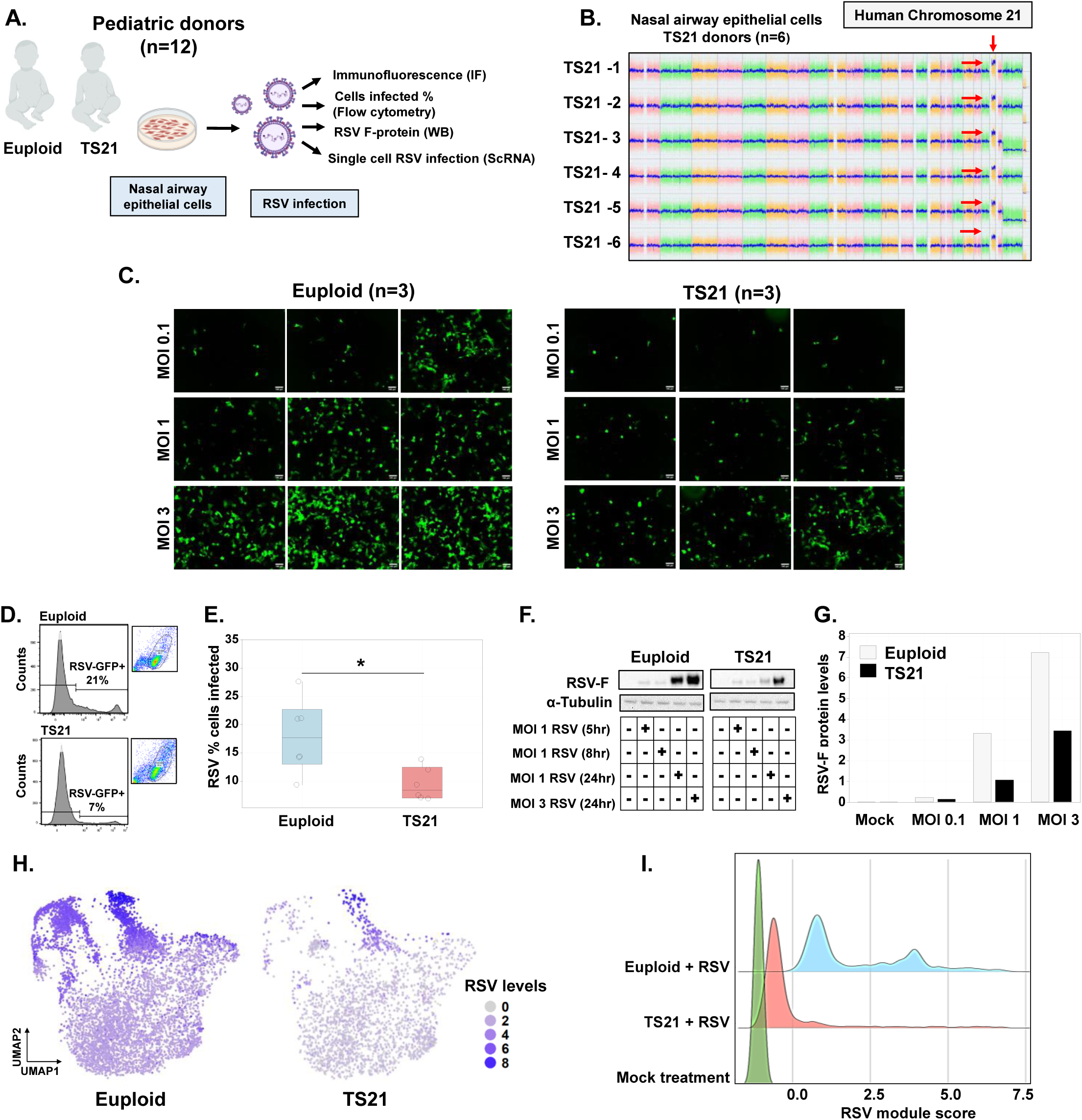
Airway epithelial cells of children with DS exhibit reduced level of RSV infection. **A.** Complementary approaches to assess RSV infection in nasal AECs. **B.** Whole genome arrays (KaryoStat+Assay) confirmed the TS21 karyotype in all primary AECs used for RSV infection assessment. **C.** Fluorescence microscopy images of GFP-RSV expression. Results representative of AECs derived from n=3 independent euploid donors and n=3 independent donors with DS. **D-E.** Flow cytometry histogram and quantification of GFP-RSV expression. Results representative of AECs derived from n=6 euploids and n=6 individuals with DS. Boxes represent interquartile ranges and medians, with notches approximating 95% confidence intervals. *p< 0.05. **F-G.** Immunoblotting and quantification of RSV F-protein expression. Results representative of AECs derived from n=6 euploids and n=6 individuals with DS. **H-I** Single-cell RNA-seq data presented as uniform manifold approximation and projection (UMAP) of GFP gene expressed by RSV (H) and ridge plots comparing a module score encompassing the ten RSV coding genes (**I**). Histogram plot showing RSV module score calculated as combined expression of 10 RSV genes detected uninfected (Mock) and RSV-infected AECs derived from an infant donor with DS (n=8,215 cells) and an age-matched euploid donor (n=14,857 cells).

To gain additional insight into the extent of RSV infection at single-cell level, we quantified RSV-encoded genes in AECs from DS and euploid donors using single-cell (sc)RNA-seq. We analyzed 23,072 cells encompassing euploid and TS21 AECs prior to and 24 hours after RSV infection (MOI 1) (**Fig. 1H-I**). Quantifying the expression of the GFP transcript expressed by RSV-GFP, we identified widespread infection of the euploid cells, with the highest level in a subset of euploid AECs (**Fig. 1H**). Using a polygenic module score that combines 10 RSV encoded genes, we identified lack of RSV gene expression in the mock treated AECs, and abundant RSV gene expression in all RSV treated AECs (**Fig. 1I**). Interestingly, while the euploid AECs show high module scores with bimodal distribution, TS21 AECs showed largely unimodal scores with RSV genes expressed below the level detected in euploid cells (**Fig. 1I**). These single cell results not only confirmed that fewer TS21 cells are infected but also provided additional evidence of reduced levels of RSV infectivity in each individual TS21 AEC compared to the euploid cells. Taken together, these findings demonstrate that TS21 AECs are not more prone to RSV infection; on the contrary, they exhibit significant protection from RSV infection.

### TS21 airway epithelial cells exhibit baseline hyperactivation of IFN signaling and desensitization to subsequent induction of JAK/STAT1 activation

Having established that TS21 AECs do not exhibit heightened RSV infectivity, and considering the severe impact of the interferonopathy of DS across various cell systems^16,18,20,27^, we next focused on defining the effects of TS21 on IFN signaling in human AECs. For this we examined the effect of TS21 on gene expression in human AECs at the single-cell level. Similar to the RSV module score, here we developed a polygenic module score for over a hundred Hsa21-encoded genes expressed in AECs (**Supplementary Table 2**). Consistent with increased gene dosage in TS21, we identified homogeneous upregulation of Hsa21 genes in all cells from the TS21 donor (**Fig. 2A**). In contrast, module scores for the IFN-regulated genes (**Supplementary Figure 1**) showed heterogeneous cell-to-cell upregulation of IFN signaling in TS21 AECs, with a distinct population of “IFN-activated cells” uniquely present in TS21 but not in euploid cells (**Fig. 2B**). IFN-activated genes overexpressed in TS21 AECs primarily include those regulated by IFN type I and III (**Supplementary Figure 2**), the type of IFNs produced by AECs^21,25^. Analyses across all cells confirmed that TS21 AECs showed overall upregulation of multiple antiviral IFN genes, including those not encoded in Hsa21 (**Fig. 2C**). Gene Ontology (GO) analysis revealed that these IFN-inducible genes represented the top hallmark pathways upregulated in TS21 AECs (**Fig. 2D**). Together, these findings demonstrate that TS21 AECs exhibit IFN hyperactivation even without a viral exposure. This is caused not merely by the increased Hsa21 gene dosage, but also due to transcriptional activation induced by IFN type I and III. To understand the dynamics of IFN I and IFN III activation in TS21 AECs, we examined JAK-STAT1 signaling, the central transcriptional pathway activated by IFN I and III^13,14^. For this we exposed AECs to either type I or type III IFN and then monitored the levels of STAT1, as well as USP18, an IFN-inducible negative regulator of JAK/STAT1 signaling that facilitates IFN receptor desensitization^28,11^. Relative to euploid AECs, we found that TS21 AECs exhibit increased baseline expression of both total and phosphorylated STAT1 (pSTAT1), as well as USP18 (**Fig. 2E-F**). In contrast, we observed a significant reduction in pSTAT1 levels in TS21 AECs in response to type I, III IFN stimulation compared to euploid cells (**Fig. 2F-G**). Taken together, these results demonstrate that children with DS are affected by an airway epithelial interferonopathy characterized by baseline hyperactivation of IFN signaling and desensitization to subsequent induction of JAK/STAT1 activation by type I, III IFNs.

**Figure 2.**
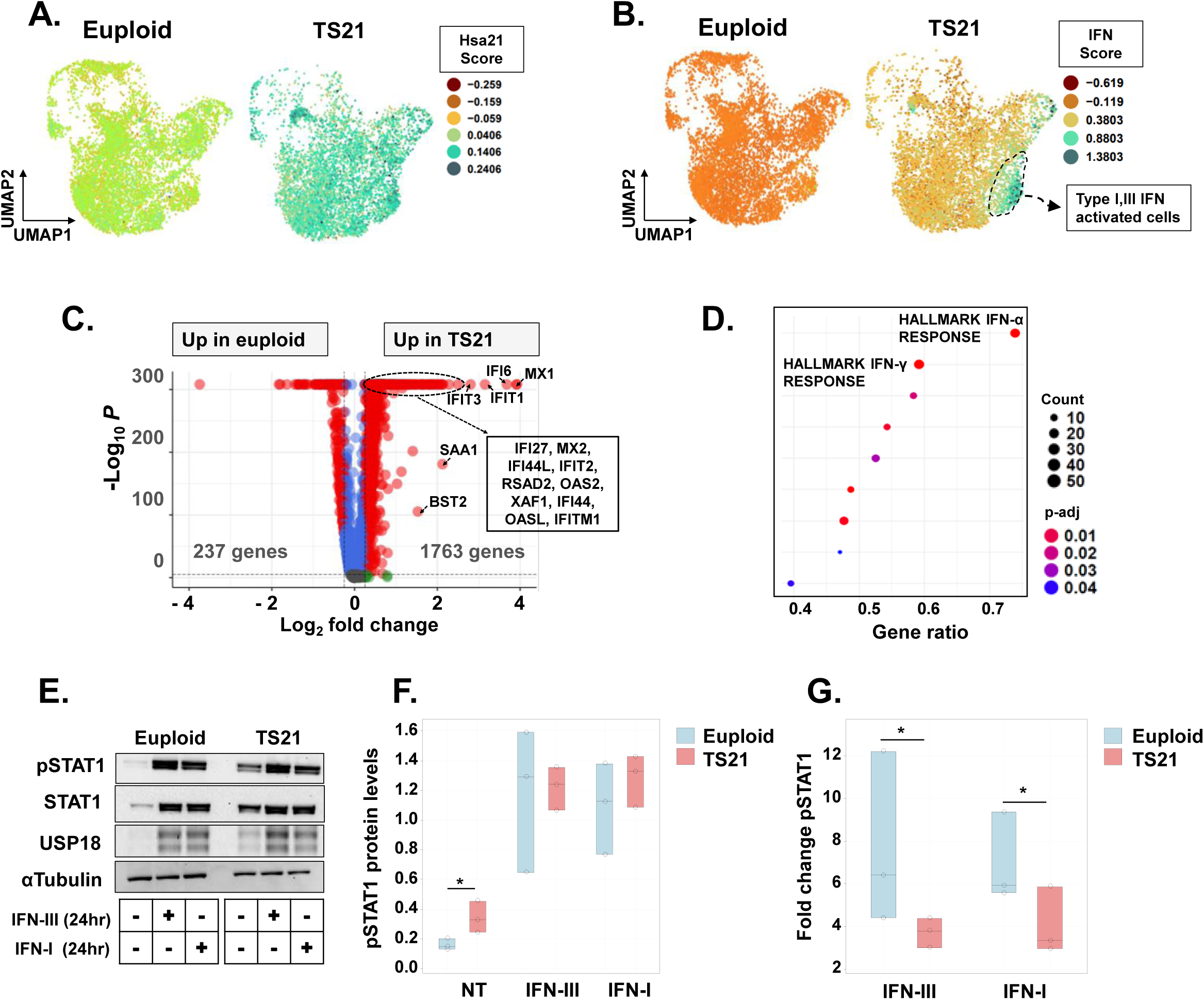
TS21 airway epithelial cells exhibit baseline hyperactivation of IFN signaling and desensitization to subsequent induction of JAK/STAT1 activation. **A-B.** UMAP of scRNA-seq data shows (**A**) homogenous overexpression of Hsa21 genes in TS21 AECs and (**B**) heterogenous overexpression of type I, III IFN-inducible genes in a subpopulation of TS21 AECs (marked in green). Results representative of uninfected AECs from an infant donor with DS (n=5,742 cells) and an age-matched euploid donor (n=7,879 cells). **C.** Volcano plot from the same dataset demonstrate multiple IFN-induced genes upregulated in TS21 AECs relative to euploid cells. **D.** Gene Ontology analysis identified IFN-activated genes as top hallmark pathways upregulated in TS21 AECs. **E.** Immunoblotting of pSTAT1, STAT1 and USP18 protein levels at baseline and after stimulation with type-III IFN/λ1 and type-I IFN/β. Results representative of AECs derived from 3 independent euploid donors and 3 independent donors with DS. **F-G.** Comparison of STAT1 normalized protein levels at baseline (**F**) and fold-changes after IFN I, III stimulation (**G**). Boxes represent interquartile ranges and medians, with notches approximating 95% confidence intervals. Results representative of AECs derived from 3 independent euploid donors and 3 independent donors with DS. *p<0.05

### Airway epithelial cells of children with DS demonstrate impaired RSV-induced IFN responses and reduced production of type III IFN

We next investigated the impact of TS21 on the induction of IFN responses in AECs during RSV infection. Consistent with our previous findings, we observed elevated basal levels of STAT1 protein in TS21 AECs but reduced induction during RSV infection compared to euploid cells (**Fig. 3A**). Similarly, MX1, a potent IFN-inducible antiviral protein encoded in Hsa21^14,29,30^, showed baseline elevation in TS21 AECs, but this failed to increase further during RSV infection, contrasting with the prominent virus-induced upregulation seen in euploid cells (**Fig. 3B**). To evaluate the impact of TS21 on the transcriptomic induction of IFN responses, we used polygenic IFN module scores to quantify IFN signaling activation at single cell level with and without RSV treatment (**Fig. 3C**). This revealed that RSV exposure of the euploid AECs resulted in an increase of over 85% in the proportion of cells with activated IFN type I, III response (**Fig. 3D**). In contrast, only 14% of TS21 AECs exhibited increased IFN type I, III response upon RSV exposure (**Fig. 3D**). These results demonstrate that TS21 is associated with increased basal, but reduced RSV-induced airway epithelial IFN responses during viral infection.

**Figure 3.**
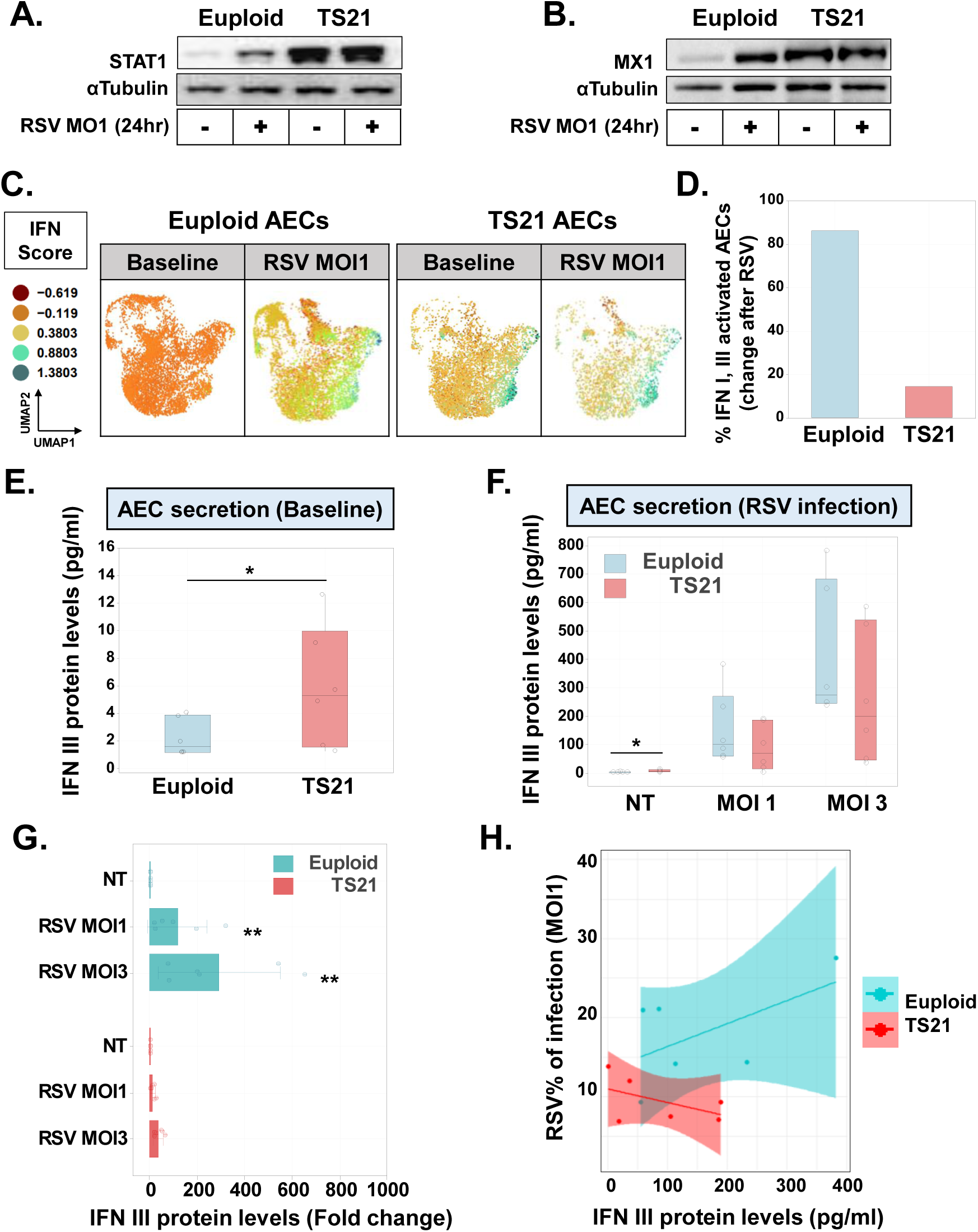
Airway epithelial cells of children with DS demonstrate impaired RSV-induced IFN responses and reduced production of type III IFN. **A-B.** Immunoblotting of STAT1 (**A**) and MX1 protein levels (**B**) at baseline and after RSV infection. Result representative of AECs derived from 3 independent euploid donors and 3 independent donors with DS. **C-D** UMAP visualization of AEC scRNA-seq data (**C**) demonstrating cells activated by type I-III IFN (marked in green-yellow) after RSV infection (MOI 1 x24 hours). (**D**) Bars representing the % of cells activated after RSV in TS21 and euploid cells (**D**). **E-G** Comparison of type-III IFN/λ1 protein levels at baseline (**E**) and during RSV infection (**F**, MOI 1 and MOI 3 x24 hours). Boxes represent interquartile ranges and medians, with notches approximating 95% confidence intervals. (**G**) Bars represent corresponding average fold changes and 95% confidence intervals relative to baseline type-III IFN levels (**G**). Results representative of AECs derived from 6 independent euploid donors and 6 independent donors with DS. **p <0.01, *p<0.05 **H.** Scattered plot showing the relationship between type-III IFN levels, and the percentage of RSV-infected cells (MOI 1 x24 hours) quantified by flow cytometry. The colored shaded area represents the 95% confidence interval around the regression line for each group. Results representative of AECs derived from 6 independent euploid donors and 6 independent donors with DS.

We then examined whether the reduced induction of IFN responses in TS21 AECs results in deficient production of airway IFN during viral infection. We focused on type III IFN since AECs do not elicit robust type I IFN responses against this virus^24–26,31^ (**Supplementary Figure 3**).

Prior to RSV infection, we observed higher levels of secreted type-III IFN protein in AECs from children with DS (**Fig. 3E**), consistent with their baseline IFN hyperactivation. However, following RSV infection, the induction of type-III IFN secretion was lower in AECs from children with DS compared to euploid children (**Fig. 3F**). When examining the fold changes relative to baseline type-III IFN levels, we noted that even when exposed to a higher viral dose (MOI 3), the suppressed secretion of type-III IFN persisted in TS21 AECs relative to that in euploid children (**Fig. 3G**). Furthermore, unlike the responses in euploid donors, AECs from children with DS failed to increase the production of type-III IFN proportionally to the number of RSV-infected cells. (**Fig. 3H**). Taken together, these findings demonstrate that children with DS have a previously unrecognized airway epithelial interferonopathy characterized by baseline IFN hyperactivation and deficient production of type-III IFN during viral infection.

### Type III IFN response deficit in TS21 airway epithelial cells is mediated by hyperactivation of JAK-STAT signaling

We next investigated the mechanism underlying reduced type III IFN responses in TS21 AECs during RSV infection. We hypothesized that the baseline activation of JAK-STAT signaling in TS21 AECs impairs subsequent signaling activation, leading to reduced virus-induced production of type III IFN. To test this hypothesis, we employed the JAK inhibitor tofacitinib, which has been used to mitigate the effects of IFN hyperactivation in individuals with DS.^20,32,33^. Our initial experiments identified that tofacitinib reversed baseline STAT1 activation in TS21 AECs in a dose-dependent manner, with effects detectable within three hours of treatment (**Fig. 4A** and **Fig. 4B**). We then examined the effect of tofacitinib on IFN signaling activation during RSV infection. Since JAK-STAT inhibition can potentially suppress the antiviral response^34^, we conducted tofacitinib experiments using two methods: continuous treatment for 24 hours over the course of RSV infection, and prophylactic treatment for only three hours, removing the drug prior to RSV infection to allow antiviral IFN signaling induction. Continuous and prophylactic tofacitinib treatments reduced baseline levels of IFN signaling proteins in TS21 AECs (**Fig. 4C**). Continuous tofacitinib treatment also inhibited IFN signaling activation during RSV infection in euploid and TS21 AECs (**Fig. 4C**). In contrast, prophylactic treatment three hours prior to virus exposure did not lead to suppression of antiviral IFN signaling (**Fig. 4C**) nor did this increase the level of RSV infection (**Fig. 4D**). These results demonstrate that prophylactic treatment with tofacitinib reverses baseline IFN hyperactivation in TS21 AECs while maintaining the protective induction of antiviral IFN signaling during RSV infection.

**Figure 4.**
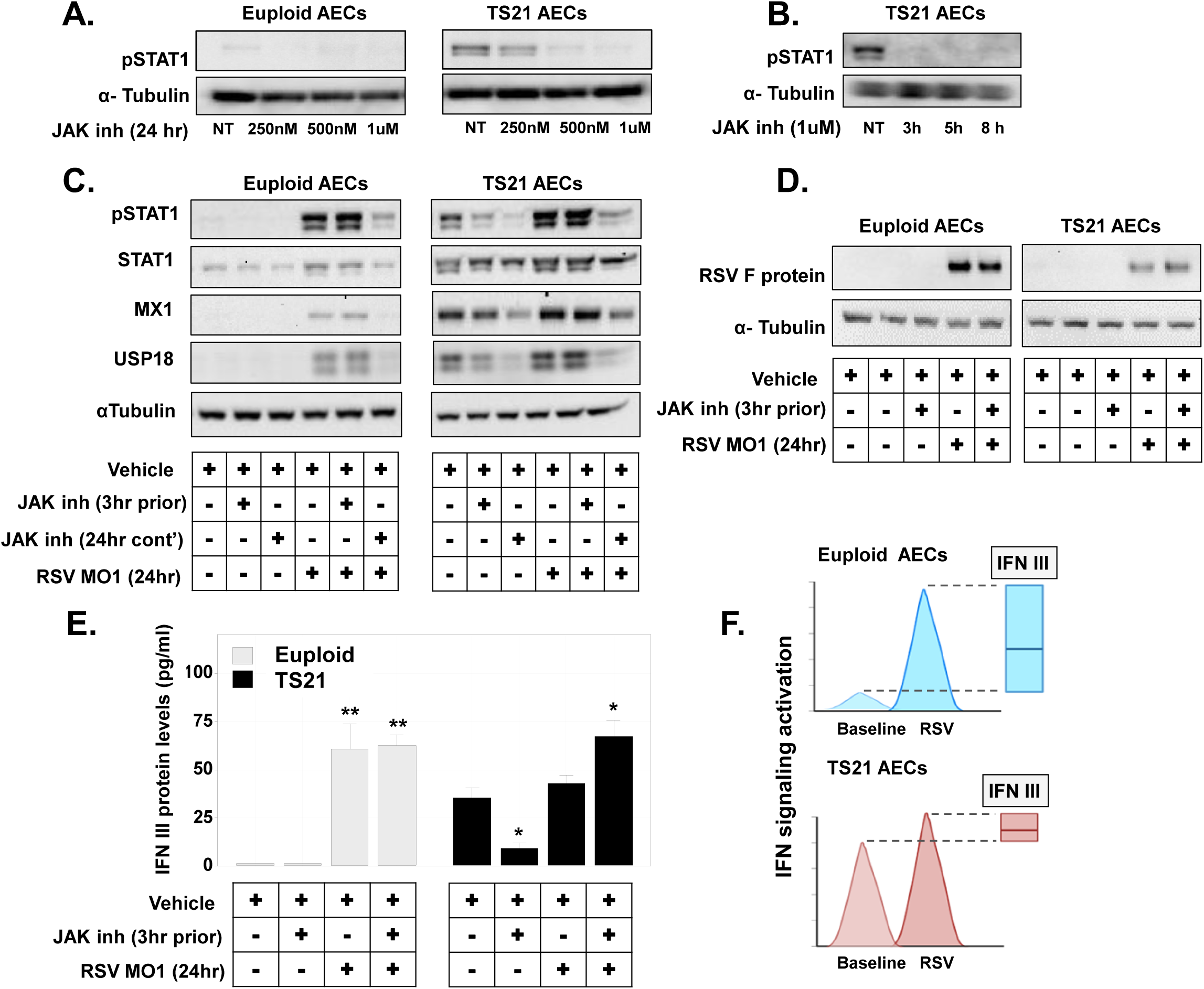
Type III IFN response deficit in TS21 airway epithelial cells is mediated by hyperactivation of JAK-STAT signaling. **A-B.** Immunoblotting of pSTAT1 protein levels after JAK inhibitor tofacitinib treatment (250nM - 1uM) (**A**) and time-response effect (**B**) in euploid and TS21 AECs. **C.** Immunoblotting of pSTAT1 and STAT1 protein levels at baseline and following RSV infection (MOI 1 x 24 hours). Data obtained before and after continuous tofacitinib treatment (1uM x 24 hours) or prophylactic treatment for three hours prior to RSV infection. **D.** Immunoblotting of RSV protein F after viral infection (MOI 1 x 24 hours) with and without prophylactic JAK inhibitor tofacitinib treatment in euploid and TS21 AECs. **E.** Bars represent average fold changes and corresponding 95% confidence intervals of type-III IFN/λ1 protein levels at baseline and following RSV infection (MOI 1 x 24 hours) before and after JAK inhibitor tofacitinib treatment. Results representative of n=3 independent experiments. Bars represent the mean ± SEM. **p <0.01, *p<0.05. **F.** Summary diagram of the airway epithelial interferonopathy in DS and impact during RSV infection.

Building on these findings, we then investigated whether inhibiting the baseline activation of JAK-STAT signaling could restore type-III IFN production during RSV infection in TS21 AECs. Consistent with our previous data, we observed that RSV elicited a robust production of type-III IFN in euploid AECs, but not in TS21 AECs (**Fig. 4E**). Prophylactic treatment with tofacitinib for three hours had no effect on type-III IFN production in euploid cells, neither at baseline nor during RSV infection (**Fig. 4E**). In contrast, we observed that in TS21 AECs, prophylactic tofacitinib treatment had a dual effect, reducing type-III IFN production while also increasing the response following RSV exposure to levels comparable to those seen in euploid cells (**Fig. 4E**). Collectively, these results demonstrate that hyperactivation of JAK-STAT signaling alters AEC antiviral immunity in children with DS, leading to a reduced capacity to mount protective type-III IFN responses during RSV infections (**Fig. 4F**).

### TS21 airway epithelial cells exhibit a dysregulated proinflammatory response during RSV infection

In addition to generating protective type III IFN responses, AECs must regulate RSV-induced inflammation to prevent severe respiratory disease caused by excessive cell infiltration and intraluminal airway obstruction^35^. Given the disproportionally high risk of children with DS to develop severe RSV-induced respiratory disease^8^, we next investigated whether alterations of the antiviral response in TS21 AECs also entail a potentially harmful proinflammatory state. For this we examined the effect of TS21 on gene expression in AECs following RSV infection (MOI 1 x 24 hours). Analyses of AEC transcriptomic profiles of children with and without DS (n=12) identified 591 differentially expressed genes after viral infection (**Fig. 5A-B**). In contrast to our findings at baseline (**Fig. 2**), following RSV infection, only 11 of the 395 upregulated genes in TS21 AECs were encoded in Hsa21 and none was linked to IFN signaling (**Fig. 5C**). GO analyses revealed that most biological processes upregulated in DS after RSV infection are related to immunity, with the notable exception of antiviral IFN responses (**Fig. 5D**). The top GO process upregulated in RSV-infected TS21 AECs was “leukocyte migration involved in the inflammatory response” (Fold enrichment 12.29, adjusted p = 1.88E-02, **Fig. 5D**). Other GO terms included “antibacterial humoral response” and “neutrophil migration” (**Fig. 5D**). The genes implicated in these processes comprised leukocyte/granulocyte chemotactic factors (e.g., CXCL5, CX3CL1, **Fig. 5E-F**), antimicrobial proteins (e.g., S100A4/7/8/9, **Fig. 5E-F**), and ALOX5, which catalyzes the biosynthesis of leukotrienes in the airways^36^. Other upregulated proinflammatory genes included the Interleukin 1 receptor (IL1R1), and the toll-like receptor 4 (TLR4), which primarily orchestrates antibacterial immunity via lipopolysaccharides recognition ^37^(**Fig. 5G**). Taken together, our results indicate that children with DS have a dysregulated AEC immune program, resulting in a paradoxical antiviral response. TS21 AECs overexpress IFN antiviral genes before viral exposure; however, during RSV infection, unlike euploid AECs, TS21 AECs preferentially upregulate pathways involved in antibacterial defense, leukocyte/neutrophil migration, and airway inflammatory responses.

**Figure 5.**
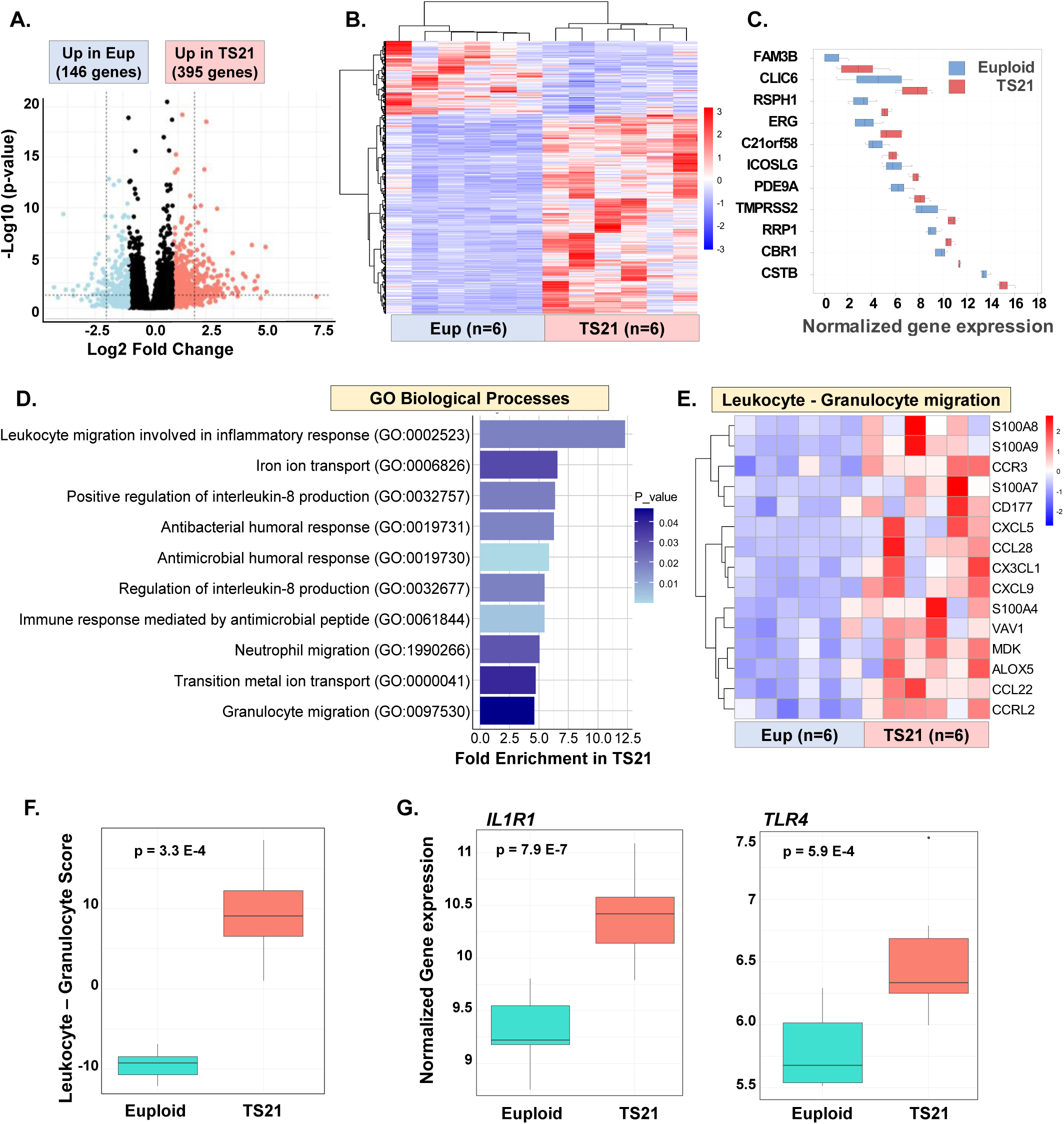
TS21 airway epithelial cells exhibit a dysregulated proinflammatory response during RSV infection. **A-B.** (**A**) Volcano plot (**B**) and heatmap showing all genes with differential gene expression (Fold Log2 Change >1 and Benjamini–Hochberg adjusted p-values <0.05) in TS21 vs. euploid children after RSV infection (MOI 1 x 24 hours). **C.** Boxplots presenting significantly overexpressed Hsa21 genes in TS21 AECs after RSV infection. **D.** Gene Ontology analysis of the same dataset showing top 10 upregulated biological processes in TS21 AECs. **E-G.** (**E**) Heatmap and (**F**) corresponding boxplot showing a polygenic score integrating the top 15 leukocyte/granulocyte migration genes with significant upregulation in TS21 AECs (Benjamini–Hochberg adjusted p-values <0.05). **(G)** Comparison of IL1R1 and TLR4 expression in TS21 vs. euploid AECs. Boxes represent interquartile ranges and medians, with notches approximating 95% confidence intervals. Data representative of AECs from n=6 euploids and n=6 individuals with DS.

### TS21 airway epithelial cells demonstrate heightened proinflammatory chemokine secretion at baseline and during RSV infection

The dysregulated immune program of TS21 AECs exposed to RSV suggests these cells exhibit a proinflammatory phenotype. To examine this possibility, we performed a cytokine array to define the secretory immune profile of AECs from children with and without DS (n=12, **Fig. 6A-B**). We included 20 analytes covering key cytokines implicated in the activation and chemotaxis of major immune cell populations to the airways (**Fig. 6B**). Using a multivariate correlation matrix integrating all 20 analytes, we identified that children with DS have an overall distinct immune AEC profile at baseline (**Fig. 6C**). Hierarchical clustering of individual cytokine values revealed that TS21 cells exhibited an exaggerated production of CXCL10 (>10-fold increase relative to euploid cells) and CXCL5 (>5-fold increase relative to euploid cells) (**Fig. 6D-E**). IL1β levels were increased 4-fold in TS21 AECs (median 0.6pg/ml in euploid vs. 2.3pg/ml in DS, p=0.004) along with other pro-inflammatory cytokines that also trended to have higher levels in TS21 AECs (i.e., CCL2, **Fig. 6D**). These data demonstrate the aberrant proinflammatory phenotype of TS21 AECs even in the absence of viral infection.

**Figure 6.**
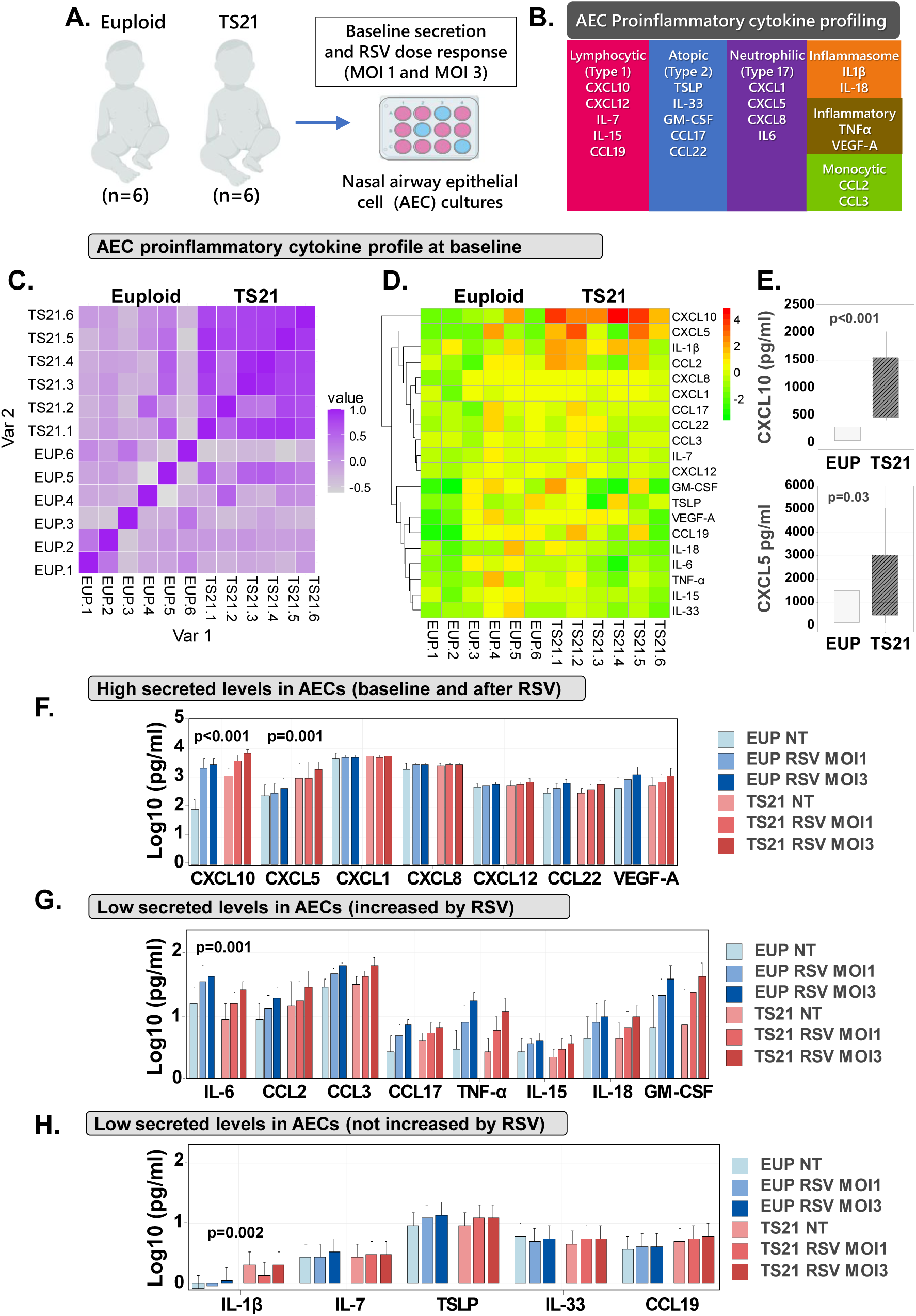
TS21 airway epithelial cells demonstrate heightened proinflammatory chemokine secretion at baseline and during RSV infection. **A-B.** (**A**) Schematic of study groups, experimental approach, and (**B**) 20 analytes included with corresponding functional immune roles. **C-D.** (**C**) Multivariate Spearman correlation matrix integrating baseline values of all 20 analytes and (**D**) unsupervised hierarchical clustering of individual cytokines from the same dataset. Data expressed as the log2-transformed fold change over the mean control (euploid) per cytokine. **E.** Boxes represent interquartile ranges and medians, with notches approximating 95% confidence comparing protein levels of CXCL10 and CXCL5, the analytes with the most significant differences at baseline in TS21 AECs (n=6) relative to the euploid group (n=6). **F-H.** Bars and corresponding 95% confidence interval means of each of the analytes at baseline and following RSV infection with MOI 1 or MOI 3 during 24 hours in euploid AECs (blue scale) TS21 AECs (red scale). Data expressed as Log10 values and grouped as analytes with high levels (**F**) and those with low levels (**G-H**), with or without significant response to RSV. Data representative of AECs from n=6 euploids and n=6 individuals with DS.

We next examined whether the proinflammatory secretion by TS21 AECs is further exacerbated by RSV infection. Monitoring the secretion of all 20 cytokines after increasing doses of RSV (MOI1 and MOI3) in euploid and TS21 cells revealed that some proinflammatory mediators are released at high levels (**Fig. 6F**), others have low levels even after exposure to a high RSV dose (**Fig. 6G-H**). Among the proinflammatory cytokines with high levels, CXCL10 and CXCL5 demonstrated a robust dose-response increase following RSV infection (≈10 Fold), which was significantly higher in TS21 AECs than in euploid AECs (**Fig. 6F**). Cytokines with low levels showed similar responses to RSV in both groups except for IL-6, which was increased in euploid cells (**Fig. 6G**), and IL1β, which showed higher levels in TS21 AECs but was not induced after increasing doses of RSV infection (**Fig. 6H**). Collectively, these results demonstrate a dysregulated AEC antiviral response in children with DS, characterized by heightened baseline secretion of proinflammatory chemokines (CXCL10 and CXCL5) that is further exacerbated during RSV infection.

### Children with DS demonstrate aberrant airway antiviral IFN III response and increased proinflammatory chemokine production during severe viral respiratory infection

Having established that AEC immune dysregulation in DS not only affects IFN signaling, but also alters the pro-inflammatory response, we next investigated the interplay between these responses during RSV infection. We first examined the relationship between the release of type III IFN and proinflammatory chemokines CXCL5 and CXCL10 in euploid and TS21 AECs exposed to RSV (**Fig 7A**). Our results demonstrated no correlation between the virus-induced production of type III IFN and CXCL5 or CXCL10 in AECs from euploid children (**Fig 7B**). This suggests that the normal protective antiviral type III IFN responses mounted in AECs limits RSV infection without concomitant CXCL5 or CXCL10 production. In contrast, in TS21 AECs, the release of type III IFN showed a strong linear correlation with the increase in CXCL5 and CXCL10 release (**Fig 7C**). Together, these findings demonstrate that unlike euploid AECs, which limit the progression of viral infections through local type III IFN antiviral defense^24–26^, immune dysregulation in TS21 AECs results in an aberrant antiviral response characterized by excessive release of proinflammatory mediators known to enhance the chemotaxis of neutrophils and other immune cells to the epithelial barrier^22,23^.

**Figure 7.**
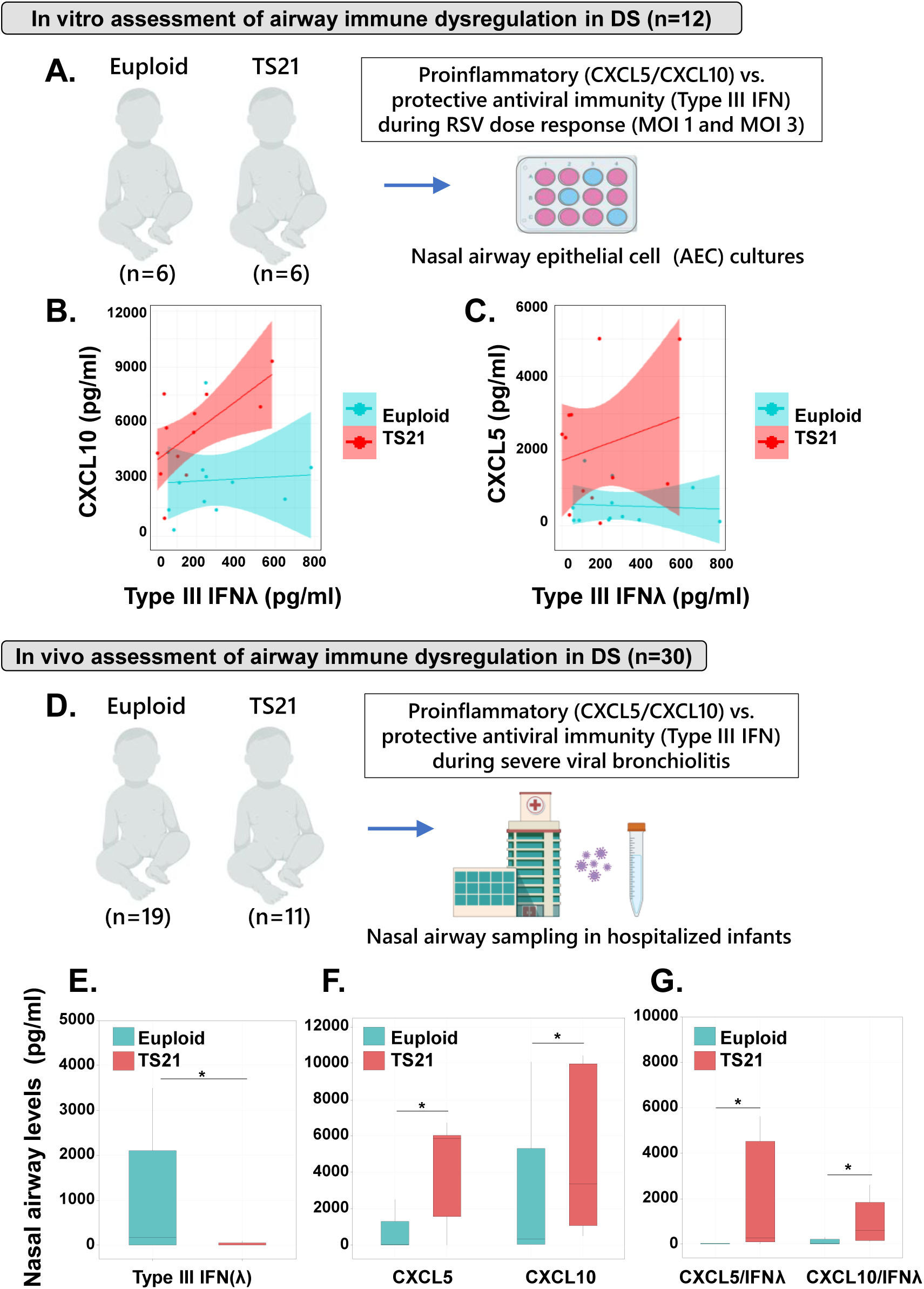
Children with DS demonstrate aberrant airway antiviral IFN III response and increased proinflammatory chemokine production during severe viral respiratory infection. **A.** Schematic of study groups and in vitro experimental approach. **B-C.** Scattered plots showing the relationship between (B) CXCL10 or (C) CXCL5 protein levels and corresponding type III IFN response in AECs exposed to RSV - TS21 cells (red) vs. euploid cells (blue). The colored shaded area represents the 95% confidence interval around the regression line for each group. Results representative of AECs derived from 6 independent euploid donors and 6 independent donors with DS exposed to RSV for 24 hours at MOIs of 1 or 3. **D.** Schematic of study groups and in vivo experimental approach. **E-G.** Boxes represent interquartile ranges and medians, with notches approximating 95% confidence intervals comparing nasal protein levels of (**E**) type III IFN, (**F**) CXCL10 and CXCL5, as well as (**G**) the ratios of CXCL5 and CXCL10 to type III IFN responses in children with DS (red, n=11) and children without DS (blue, n=19). *p<0.05.

Based on these in vitro findings, we hypothesized that while baseline IFN hyperactivation in TS21 confers initial protection against viral entry into AECs, in the setting of advanced severe viral respiratory illnesses requiring hospitalization, children with DS are mostly affected by the proinflammatory effects of dysregulated airway antiviral immunity. To test this postulate, we analyzed a group of infants and young children hospitalized due to severe viral bronchiolitis caused by RSV or other respiratory viruses (n=30, **Fig 7D**. **Supplementary Table 1**). (**Fig. 7E**). Comparison of the levels of type III IFN, CXCL5, and CXCL10 in the nasal mucosa of children with DS (n=11) relative to age- and sex-matched euploid children (n=19) showed an impaired airway IFN response in DS during severe viral respiratory infections, with a significant reduction in type III IFN secretion (median euploid= 176 pg/ml vs. DS= 3.6 pg/ml, p= 0.01) **(Fig. 7E).**

Similar to our observations in AECs, severe viral respiratory illnesses led to an increased airway proinflammatory response in children with DS as compared to their euploid counterparts, with exaggerated production of CXCL5 (median euploid: 18 pg/ml vs. DS: 5832 pg/ml, p= 0.04, **Fig 7F**) and CXCL10 (median euploid: 353 pg/ml vs. DS: 3372 pg/ml, p= 0.04, **Fig 7F**). Comparison of the ratios of CXCL5 and CXCL10 to type III IFN responses in the same individuals, identified a marked dysregulated airway immunity in children with DS, favoring proinflammatory changes over the generation of protective antiviral IFN responses. (**Fig 7G**). Therefore, in the clinically relevant context of severe viral respiratory illnesses requiring hospitalization, children with DS exhibited an imbalanced airway immune response characterized by reduced local type III IFN production and disproportionately heightened release of airway proinflammatory chemokines.

## DISCUSSION

DS is associated with multiple conditions with higher occurrence rates across all age groups^1^. Among the pediatric population, respiratory infections^2–7^, particularly those caused by RSV^8,9^, are the leading cause of hospitalization in children with DS^2–7^. As the impact of TS21 on human airway antiviral responses is largely unknown, it remains unclear why children with DS have a nine-fold increase in mortality risk from RSV compared to those without DS^8^. It also remains unexplored whether this is due to heightened viral infectivity in AECs or the immune dysregulation typical of DS^10–12^. Our study addresses this issue by demonstrating that AECs in children with DS exhibit a previously unrecognized immune dysregulation state that manifests as interferonopathy with elevated JAK/STAT1 signaling at baseline and impaired IFN activation during RSV infection. This altered antiviral response by AECs notably reduces the production of type-III IFN during RSV exposure, hampering this crucial response for respiratory mucosal antiviral immunity^24–26^. Furthermore, AECs from children with DS exhibit an exaggerated proinflammatory response at baseline and during RSV infection. This response is characterized by enhanced production of chemokines associated with leukocyte/neutrophil trafficking and virus-induced airway inflammation (CXCL5 and CXCL10)^22,23^. Thus, our study supports the potential role of modulating dysregulated AEC immune responses as a novel strategy to prevent severe viral respiratory illnesses in people with DS.

Several studies have focused on defining the dysregulated IFN responses in DS across multiple immune compartments^16–19^, but the status of these responses in AECs is lacking. Here we focused on type-III IFN, also known as IFN-λ, as it provides essential antiviral roles within the AECs^24–26^. This molecule is primarily produced by epithelial cells^24–26^, offering localized protection at the site of viral entry and replication. Although IFNs have overlapping functions^24–26^, type-III IFN is the earliest and predominant IFN produced by AECs during RSV and other viral respiratory infections ^24–26,31^. Unlike type-I or type-II IFNs, type-III IFN supports potent antiviral functions with the unique feature of not eliciting proinflammatory responses^24–26^.

Notably, the type-III IFN receptor is also primarily expressed in epithelial cells, supporting an specialized local role in protecting mucosal surfaces without triggering systemic inflammation^24–26^. This local antiviral function of type-III IFN is particularly relevant for individuals with DS, considering their systemic IFN hyperactivation and proinflammatory state^10–12^, which could be exacerbated by the failure to eliminate viruses at the point of entry leading to global overactivation of IFN and cytokine responses^38^.

An important finding of our study is that the baseline IFN hyperactivation of TS21 AECs is associated with reduced RSV infectivity in vitro. This finding aligns with observations from over 40 years ago^39^, indicating heightened viral resistance in TS21 fibroblasts^39^. Viral resistance is expected in TS21 cells as they have increased baseline expression of Hsa21-encoded IFN-inducible antiviral effectors including MX1 and MX2^14,29,30^. Importantly, despite being associated with reduced viral infectivity, the interferonopathy of TS21 AECs leads to deleterious effects following RSV exposure. This includes impaired activation of IFN signaling and deficient virus-induced production of type-III IFN in children with DS. This finding is clinically relevant given the strong evidence linking suppressed local AEC type-III IFN responses to worse respiratory outcomes in infants with viral bronchiolitis^40–42^. Studies involving nasal sampling of infants with RSV bronchiolitis have identified that those with severe respiratory disease have significantly reduced airway type-III IFN levels and, unexpectedly, lower viral loads than those with mild RSV disease^40–42^. These observations suggest that the suppressed induction of the antiviral type-III IFN response in TS21 AECs likely has a greater clinical impact than the reduced viral infectivity seen in these cells.

In addition to reduced protective local type III IFN responses, we also identified that TS21 AECs exhibit a proinflammatory state characterized by enhanced secretion of CXCL5 and CXCL10, resembling the systemic cytokinopathy of DS^16–19^. Importantly, our AEC cytokine profiling demonstrates that the proinflammatory abnormalities caused by TS21 are cell-specific, as several cytokines with elevated systemic levels in DS did not show increased production in TS21 AECs. For instance, IL-6, which is upregulated in immune cell populations in DS ^16–19^, showed reduced levels at baseline and after RSV infection in TS21 AECs. On the other hand, despite only a few reports linking DS to inflammasome activation^43^, TS21 AECs exhibited increased baseline secretion of IL-1β. These novel findings in TS21 AECs strongly support the necessity of reporting the diverse immunological effects of TS21 across tissue-resident cell populations, particularly within the respiratory system, given the significant impact of respiratory disorders in individuals with DS^2–5^.

We also identified that the AEC dysregulation of DS is coupled to a unique transcriptomic program. At baseline, TS21 cells predominantly overexpress IFN signaling genes resembling the transcriptional signatures observed in response to viruses. Conversely, following viral infection, these IFN signatures are no longer predominant, and transcriptomic changes reflect overexpression of leukocyte/neutrophil chemotactic factors that promote airway inflammation and antibacterial defense. This pattern of dysregulated AEC antiviral response has been linked to clinical severity during RSV infections^44,45^. Experimental RSV infection in humans and mice has identified that mucosal overexpression of leukocyte/neutrophil chemotactic factors is highly predictive of severe RSV disease^44^, coinciding with reports of severe airway neutrophilic infiltration in children suffering from severe RSV infections^45^. Given the clinical relevance of our findings, future studies must investigate whether the shift from antiviral IFN signaling to a pro-inflammatory program during RSV infection in TS21 AECs is solely related to impaired IFN signaling activation, or if there are additional potentially targetable proinflammatory pathways contributing to dysregulated AEC antiviral immunity in DS.

A key conclusion of our study is that severe viral respiratory illnesses in children with DS are not due to impaired control of viral infection in AECs. Instead, we found that children with DS suffering from severe viral bronchiolitis display a dysregulated airway immunity characterized by reduced protective type-III IFN responses and excessive production of proinflammatory mediators. Although TS21 cells may initially be protected against viral infection, in the context of an advanced respiratory disease state requiring hospitalization, children with DS are predominantly affected by inflammatory changes post-infection induced by chemokines such as CXCL5 and CXCL10.These chemokines contribute to early immune cell recruitment and activation in the airways, but their sustained dysregulated production exacerbate inflammation and tissue damage in several respiratory conditions^22,23^. The excessive production of CXCL5, also known as epithelial-derived neutrophil-activating peptide 78 (ENA-78), can lead to neutrophil-mediated airway injury during respiratory infections^22^. Similarly, long-term exposure of chemokine CXCL10 causes bronchiolitis-like inflammation with severe peribronchiolar and perivascular lymphocyte infiltration^23^. Therefore, our study not only sheds light on the potential mechanism underlying severe viral respiratory illnesses in children with DS, but also provides novel human-based data supporting that targeting dysregulated airway immune responses in people with DS is a promising therapeutic opportunity to mitigate the high respiratory morbidity and lethal impact of RSV in this vulnerable population.

## Limitations of the study

We recognize that additional insights into impaired antiviral responses in DS could be gained through investigations involving differentiated AECs and fresh airway specimens from virally infected individuals with DS. Research involving a larger cohort of individuals with DS including IFN and proinflammatory profiling across multiple immune compartments can offer a more comprehensive understanding of deficits in cell and antibody-mediated immunity^18^. Such investigations are necessary to fully elucidate the mechanisms mediating the high morbidity and mortality during RSV infections in children with DS.

## Acknowledgements

We thank all families and children with and without DS for participating in this project. We are also grateful for the technical expertise provided by The Genomics Core at The Research Institute of Children’s National Hospital in Washington D.C., as well as The Single Cell & Transcriptomics Core and The Genetic Resources Core Facility (GRCF) at Johns Hopkins University. Special thanks to the Leadership Teams of the National Institutes of Health (NIH) INCLUDE (INvestigation of Co-occurring conditions across the Lifespan to Understand Down syndromE) Project and the National Lung, Heart, and Blood Institute (NHLBI) Neonatal and Pediatric Lung Disease/Critical Care Program.

This work was supported by NIH grant R01HL159595 (G.N. and J.K.J.); NIH grant R01HL141237 (G.N.); NIH grant K23HD104933 (M.J.G.); and NIH grant R01HL153604 (J.L.G.).

## Author contributions

E.C., S.B., J.K.J., G.N. contributed to conceptualization, methodology, investigation, and writing. E.C., S.B., B.S.B., A.W., K.I. conducted the experiments. S.B., H.M., M.J.G., J.L.G., E.C., J.K.J., G.N. contributed to data analysis, and visualization. C.M.L., D.P., D.K.P., M.J.G., G.N. contributed to subject enrollment, sample collection, methodology, investigation, and writing.

J.K.J. and G.N. contributed to conceptualization, supervision, funding acquisition, project administration, and writing—original draft, review, and editing.

## METHODS

### Cell collection, isolation, and culture

This study was approved by the Institutional Review Board (IRB) at Children’s National Hospital (CNH) (IRB Pro00012323, Pro00003441, STUDY00000511) and included parental written informed consent. All patient data were de-identified. Demographic information of recruited patients is provided in **Supplemental information**. Nasal airway secretions were collected at the onset of acute respiratory illnesses by a standard nasal lavage technique with 3–4 mL of sterile normal saline and immediately frozen at -80°C until time of assay^50–52^. Nasal AECs were obtained using 2.7 mm cytology brushes rotated in each nostril. Brushes with cell samples were immediately placed in transport media, airway epithelial growth media (PromoCell) and left on ice until cells could be isolated from the brush. Cells were spun down at 500g for 5 minutes at 4°C and were immediately resuspended in Conditional Reprogrammed Cell (CRC) media^50–52^. The cell suspension was placed into a PureCol Coated T-25 flask (CellStar, VWR) and allowed to rest overnight. After overnight rest, cell culture media was removed and replaced with fresh CRC media. Cells were allowed to grow to confluency, with media changes every 2-3 days before expansion to a T-75 flask (CellStar, VWR).

### Respiratory Syncytial Virus (RSV) Stock preparation

Hep2 cells were seeded at 6 million cells per T-175 flask and allowed to rest overnight. Prior to infection, one flask of Hep2 was counted to confirm expected cell numbers and allow accurate calculation of viral load. RSV-A2 GFP tagged was diluted in serum free EMEM (ATCC) media. Hep2 cells were exposed to 5mL of diluted virus stock for a final concentration of MOI 0.1 for 2 hours with occasional rocking. After 2 hours, 6mL of EMEM supplemented with fetal bovine serum (FBS, ATCC) were added to each flask. Virus was allowed to propagate for 96 hours, or until greater than 50% CPE was seen before harvesting. At time of harvest, supernatant was pooled from all flasks and kept on ice. Cells were covered with 20% sucrose (w/v) in NT buffer and immediately placed at -80°C until the layer froze. Cells were then allowed to thaw gently on ice. Once thawed, cell lysates were scrapped using a cell lifter (VWR) and the lysate was combined with the virus containing supernatant. This crude virus containing solution was then spun at 1000xG for 10 minutes at 4°C to remove cellular debris. The clarified supernatant was then transferred to a new tube. 50% PEG6000 in NT buffer was added to the supernatant for a final concentration of 10% to precipitate virus particles. Rubes were then rocked at 4C for 2 hours, after which, they were centrifuged at 3200xG for 20 minutes at 4°C to pellet virus particles. The resulting pellet was then resuspended in approximately 20x the volume of 20% sucrose in NT buffer. Resuspended virus solution was layered onto a sucrose gradient of 60% and 30% sucrose in NT buffer. Sucrose gradient with virus supernatant tubes were loaded into an SW41Ti rotor and spun at 50,000xG for 1 hour at 4°C. Tubes were then carefully removed from the rotor and a hazy band around the 30/60% sucrose interface was observed. This band was carefully collected and immediately snap frozen in cryovials using a dry ice/ethanol bath. Aliquoted viral stocks were stored at -80°C or below until time of quantification of downstream application.

### RSV infection of human airway epithelial cells

6 well plates were coated with 1mL of PureCol solution (Advanced BioMatrix) diluted 1:30 in sterile H2O. PureCol solution was allowed to incubate in the wells for 1 hour, covered, before being removed. Wells were then washed 3 times with sterile 1x PBS (Gibco). The final wash was removed, and plates were left to dry inside the biosafety cabinet. Once dry, plates were wrapped in foil and stored at 4°C until time of experiment, but no longer than 3 weeks.

Human nasal AECs were seeded on PureCol coated plates at 150,000 cells/well in 2mL CRC media. Cells were allowed to rest for 18-24 hours before CRC media was replaced with serum-free airway epithelial growth media (referred to as Serum Free Media or SF media). SF media is composed of airway epithelial growth media (Promocell). Cells were serum starved for 18-24 hours before introduction of virus. Virus stocks were diluted in airway epithelial growth media containing only retinoic acid (RA only media) to facilitate minimum growth requirements in conditionally reprogrammed cells. Virus stocks were diluted to an MOI of 1 per well.

### Interferon (IFN) and JAK inhibitor treatments

Human AEC cultures were seeded in a 6-well plate (1.5 × 10^5 cells/well). The cells were transitioned to serum-free media 24 hours later and then treated with either IFN-λ/IL-29 (100 ng/mL) or IFN β (1 ng/mL) in serum-free media. For the JAK inhibitor experiments, after being serum-starved for 24 hours, cells were plated on 6-well collagen-coated plates and then initially exposed to various concentrations (250 nM - 1 µM) of tofacitinib (Sigma-Aldrich). After dose and time response optimization, nasal AEC seeded in 6-well plates were treated with 1 µM tofacitinib for 24 hours, including during RSV infection, or for 3 hours and then washed prior to RSV exposure.

### Cytokine quantification

Nasal AEC supernatants and nasal aspirates were assayed for quantification of cytokine protein levels. All samples were stored at -80°C until the time of assay, then gently thawed on ice and spun down at 1000g for 5 minutes. A custom U-plex plate from MesoScale Discovery (Rockville, MD) was designed to evaluate type III IFN-λ/IL-29, type I IFN-β, CXCL10, CXCL5, IL-1β, TSLP, GM-CSF, IL-6, CXCL8, CCL2, CCL22, CCL3, CCL17, TNF-α, VEGF-A, CXCL1, IL-15, IL-18, IL-33, IL-7, CCL19, CXCL12 in undiluted cell supernatant samples. We quantified in nasal aspirates type III IFN-λ/IL-29, CXCL10 and CXCL5. All samples were assayed in duplicate using the manufacturer’s protocol. Data were visualized using the Discovery Workbench software.

### Western blotting

At the time of cell harvest, wells were washed with ice-cold 1x PBS, and then 170 µl of RIPA buffer (Pierce, Thermo Fisher) with protease and phosphatase inhibitors (Roche, PhosSTOP, Halt protease inhibitor) was added to each well. Cells were immediately scraped and transferred to an Eppendorf tube. Protein lysates were then transferred to a tube rotator at 4°C for 30 minutes before being stored at -80°C until the time of protein quantification. Protein concentration was determined using a Bradford assay (Pierce, Thermo Fisher).15-30ug of protein were prepared for western blotting, with equal concentrations of protein loaded across the gel. Protein lysates were treated with reducing agent and heated to 95°C for 10 minutes prior to loading on a 4-12% bis-Tris 1.5mm mini gel (Thermo fisher). Protein samples were run using an Invitrogen gel dock at 115v for 1 hour and 45 minutes. Protein was then transferred using the BioRad Turbo Transfer system onto a nitrocellulose membrane. Membranes were blocked for 1 hour in TBST with 5% milk. After blocking, membranes were probed with primary antibody according to manufacturer’s protocol. Antibody targets include: pSTAT-1, STAT1, USP18, MX1, RSV-F, and αTubulin (detailed product information in table provided below). Membranes were probed overnight with primary antibody at 4°C with gentle rocking. The following morning, membranes were probed with corresponding secondary antibodies: anti-rabbit or anti-mouse (detailed product information provided in the table below). Membranes were incubated with secondary antibody for 1 hour at room temperature with gentle shaking. After incubation, membranes were washed 3 times with TBST and subsequently developed on the BioRad ChemiDoc using either Cytivia Amersham ECL or SuperSignal West Femto Maximum Sensitivity Substrate reagent. Western blot densitometry was performed using the ImageLab Software (BioRad version 5.2.1). All samples were normalized to a-tubulin as a loading control prior to statistical analysis.

### Flow Cytometry

At time of cell harvest, media was removed from wells and washed with 1 mL of PBS. To detach cells from plates, 0.5 mL of accutase was added to each well. Plates were placed back in the incubator at 37° C for 5 to 7 minutes to allow for detachment; plates were looked at under the microscope to see if cells have fully detached. After incubation, accutase was diluted with 1.5 mL of PBS. Using a serological pipette, the well was washed using the cell suspension solution and entire 2 mL volume was placed in a FACS tube labeled for each condition.

FACS tubes containing cells were spun down in the centrifuge at 500 G for 5 minutes. Supernatant was removed from the tubes by decanting. Cells were resuspended in 2 mL FACS buffer (PBS with 1% FBS). Cells marked for viability staining were given 5 μL of 7 AAD viability dye (ThermoFisher) just before running on the MACSQuant Analyzer. All samples were placed on ice prior to run.

Flow cytometry analysis of samples was run on MACSQuant Analyzer (Miltenyi Biotech). Forward and Side Scatter channels were set to capture events that are approximately the size and granularity of our cells. Approximately 200 µL of each sample was recorded at medium flow speed after gentle mixing. Dead cell exclusion for flow cytometric analysis with the MACSQuant Analyzer was achieved by 7 AAD staining of cells and captured in the PerCP channel set to 525V. RSV infection was detected via GFP tagging of RSV prior to infection, this was captured with the MACSQuant Analyzer in the FITC channel set to 254V. All events captured for each sample were analyzed in FlowJo and gated for viability and infection using negative and single-color controls.

### Imaging

Infection by RSV-GFP was visualized prior to downstream assays using the Leica PAULA cell imager system. GFP was visualized utilizing a green fluorescence LED and 10x objective.

### RNAseq methods

Airway epithelial cells were harvested in TRIzol reagent (ThermoFisher), and total RNA was isolated using phase separation via the phenol-chloroform method. RNA quality was assessed using the Agilent 2100 Bioanalyzer.

### Single cell sequencing of human airway epithelial cells

Single-cell RNA-seq libraries were generated using the Chromium Next GEM Single Cell v 3.1 (10x Genomics, US). The cell suspensions were diluted to a density of 1000 cells/μL in 1x PBS with 0.04% BSA. Gel Beads-in-emulsion (GEMs) were generated and loaded onto Chromium Next GEM Chip G targeting 10,000 cells. After GEM generation, the GEMs were incubated to produce barcoded full-length cDNA. The cDNA was purified using silane magnetic bead and amplified using PCR. 3’ Gene Expression Library was constructed with appropriate sample indexes following 10x protocol with final libraries containing Illumina paired-end constructs (P5 and P7) compatible with Illumina sequencer. The quality of the libraries was determined using the Agilent High Sensitivity D5000 Screen Tape. The libraries were sequenced on NovaSeq (Illumina) S2, with paired end format (28×10×10×90) as recommended by 10x Genomics.

### Bulk RNAseq differential gene expression analyses

After RNA isolation, quality control was estimated using fastqc (version 0.11.9) for individual samples and multiqc for all samples^53^. After quality trimming, the reads were aligned to the human reference (hg38), and the counts estimated with RSEM (version1.3.1)^54^. For differential expression, we first filtered out low read counts (< 200 reads) and then estimated sample quality using principal component analyses (PCA). Normalization and differential expression were performed with DESeq2 (version 1.38.3) using an adjusted p-value threshold of <0.05 and an average log2 fold change of >1.

### Single cell analysis

Pre-processing of the single-cell FASTQ samples, including demultiplexing, alignment, count, and quality visualization, was performed using Cell Ranger (version 6; 10X Genomics). For the reference, we used a concatenated genome file comprising the human genome reference (hg38) from Ensembl, RSV genome reference from GenBank, and R125-eGFP transcript sequences. The samples were aligned to this reference to not only capture the expression of RSV and R125-eGFP in the RSV-treated samples but also to check for any RSV contamination in the control samples. Next, we used the Seurat scRNA-seq pipeline for further downstream analysis on each sample individually. We filtered genes present in at least 10 cells, cells with low gene and transcript numbers (<200) and removed cells with greater than 15% mitochondrial content per sample. Next, normalization was performed on each of the samples using SCTransform with 2000 variable genes, followed by doublet removal using DoubletFinder^55^.

PCA was then performed, followed by KNN graph using the first 30 PCAs, clustering using a resolution of 0.1, and Uniform Manifold Approximation and Projection for Dimension Reduction (UMAP) for visualization. FeaturePlot function from the Seurat package was used for visualizing the UMAP.

All the samples were then integrated, followed by creation of PCA, KNN graph (first 30 PCs), clustering (using a resolution of 0.1) and UMAP. Individual pairwise integration of different conditions was performed in the same fashion for differential expression analysis. Pseudo bulk analysis, using Wilcox t-test was performed between each condition with a significance level of p-value < 0.05 and log fold change > 0.25 or <0.25. Gene ontology was performed using the Hallmark genes dataset using the fgsea package in R.

### IFN module score and IFN-activated cells quantification

To examine interferon gene activation at the single-cell level in euploid and trisomy single-cell samples before and after RSV treatment, we used the AddModuleScore function from Seurat. The module scores serve as a ‘proxy’ for the differential expression of a gene list per cell and are used for annotating gene clusters. We constructed four module scores using: 1) genes encoded chromosome 21 expressed in human AECs, 2) core IFN genes induced by all types of IFNs, 3) genes only induced by type I IFN-β and III IFN-λ representing IFNs produced by AECs, and 4) genes only induced by type II IFN-γ, which is not produced by AECs. The genes utilized to construct the module scores are reported in the **Supplemental Information**. We visualized the module score for each condition using UMAP. The module score classified AECs as IFN-activated cells using ‘0’ as the threshold value. Quantification of activated cells was expressed as the proportion of cells relative to the total number of cells sequenced in each condition.

### Trisomy 21 and RSV gene module scores

To examine the expression of human chromosome 21 genes in TS21 compared to euploids, we constructed chr21 module scores using the genes expressed in AECs (113 genes reported in **Supplemental Information**). Similarly, to evaluate the expression of RSV genes in the TS21 and the euploid samples, we made RSV gene module score using the 10 RSV coding genes (NS1, NS2, N, P, M, SH, G, F, M2, and L) and the R125-eGFP transcript. The RidgePlot function (Seurat), was used to observe the distribution of the RSV gene module score, in the RSV treated and mock treated samples.

### Statistical analysis

Statistical significances were calculated using non-parametric T test for two group comparisons and with linear regression models for time or dose responses. All data were analyzed and visualized with the Minitab Statistical Package V.19.1. (Minitab, Inc., State College, PA) and/or R studio (Version: 2023.03.1+446). P-value are denoted as: ∗p<0.05, ∗∗p<0.01. Error bars on all figures represents standard error of mean (SEM) or 95% confidence interval (CI) means. Boxplots denote median values and 25th-75th percentile of each group’s distribution of values.

