## Supplementary data for "Dysregulated airway epithelial antiviral immunity in Down Syndrome impairs type III IFN response and amplifies airway inflammation during RSV infection"

**Supplementary Table. 1. Study subjects**

| **Nasal AEC donors** |  |  |  |
| --- | --- | --- | --- |
| **Category** | **Total** | **Euploid** | **TS21** |
| **Number of subjects (n)** | 12 | 6 | 6 |
| **Mean age at enrollment, years (SD)** | 5 (4.1) | 4.7(3.9) | 5.7(4.7) |
| **Sex, male %, (n)** | 58.3 (7) | 83.3 (5) | 33.3 (2) |
| **Race/Ethnicity %, (n)** |  |  |  |
| **Caucasian** | 8.3 (1) | 0 (0) | 16.7 (1) |
| **African American** | 75 (9) | 83.3 (5) | 66.7 (4) |
| **Hispanic** | 0 (0) | 0 (0) | 0 (0) |
| **Other** | 16.7 (2) | 16.7 (1) | 16.7 (1) |
| **Nasal aspirates** |  |  |  |
| **Category** | **Total** | **Euploid** | **TS21** |
| **Number of subjects (n)** | 30 | 19 | 11 |
| **Mean age at enrollment, years (SD)** | 1.78 (1.25) | 1.75 (0.78) | 1.82 (1.8) |
| **Sex, male % (n)** | 63 (19) | 63 (12) | 63 (7) |
| **Race/Ethnicity % (n)** |  |  |  |
| **Caucasian** | 17 (5) | 11 (2) | 27 (3) |
| **African American** | 50 (15) | 58 (11) | 36 (4) |
| **Hispanic** | 30 (9) | 26 (5) | 36 (4) |
| **Other** | <1 (1) | <1 (1) | 0 (0) |
| **RSV % (n)** | 50 (15) | 63 (12) | 27 (3) |
| **Rhinovirus % (n)** | 47 (14) | 53 (10) | 36 (4) |
| **Mixed infection % (n)** | 30 (9) | 37 (7) | 18 (2) |
| **Human metapneumovirus % (n)** | 13 (4) | <1 (3) | <1 (1) |
| **Adenovirus % (n)** | 13 (4) | <1 (1) | 27 (3) |
| **Parainfluenza % (n)** | 13 (4) | <1 (1) | 27 (3) |

**Supplementary Table. 2. Hsa21-encoded genes expressed in AECs (n=113 genes)**

| CXADR | C2CD2 | ADAMTS1 | POFUT2 | ITGB2 |
| --- | --- | --- | --- | --- |
| RWDD2B | ZBTB21 | MIS18A | TMPRSS2 | KRTAP19-1 |
| MAP3K7CL | CBR1 | WDR4 | MX1 | TIAM1 |
| URB1 | CBR3 | NDUFV3 | MX2 | GATD3 |
| USP16 | RUNX1 | GABPA | LCA5L | ICOSLG |
| CYYR1 | EVA1C | CLIC6 | ATP5PF | IFNAR2 |
| MRPS6 | CFAP298 | PIGP | SPATC1L | PCNT |
| SLC5A3 | CFAP298-TCP10L | SYNJ1 | LSS | IFNAR1 |
| CHAF1B | BTG3 | RBM11 | HLCS | SON |
| SCAF4 | SUMO3 | HSPA13 | KCNE1 | RCAN1 |
| CCT8 | MRPL39 | PSMG1 | COL6A1 | ADARB1 |
| LTN1 | SMIM11 | BRWD1 | TFF3 | NRIP1 |
| APP | SIM2 | HMGN1 | USP25 |  |
| CLDN8 | RIPPLY3 | GET1 | SOD1 |  |
| HUNK | RRP1B | DSCAM | DOP1B |  |
| C21orf91 | PDXK | BACE2 | PRMT2 |  |
| MORC3 | CSTB | RIPK4 | ERG |  |
| PAXBP1 | RRP1 | ABCG1 | B3GALT5 |  |
| C21orf62 | AGPAT3 | RSPH1 | MCM3AP |  |
| ITSN1 | TRAPPC10 | SLC37A1 | YBEY |  |
| CLDN14 | PWP2 | PDE9A | C21orf58 |  |
| KCNJ6 | DYRK1A | SIK1 | DIP2A |  |
| ATP5PO | PRDM15 | HSF2BP | DNAJC28 |  |
| CHODL | PKNOX1 | H2BC12L | GART |  |
| BACH1 | CBS | PFKL | DONSON |  |
| ETS2 | U2AF1 | CFAP410 | TMEM50B |  |
| KCNJ15 | SETD4 | LRRC3 | IL10RB |  |
| CRYZL1 | SLC19A1 | UBE2G2 | IFNGR2 |  |
| N6AMT1 | COL18A1 | PTTG1IP | VPS26C |  |
| ADAMTS5 | FAM3B | SLX9 | TTC3 |  |

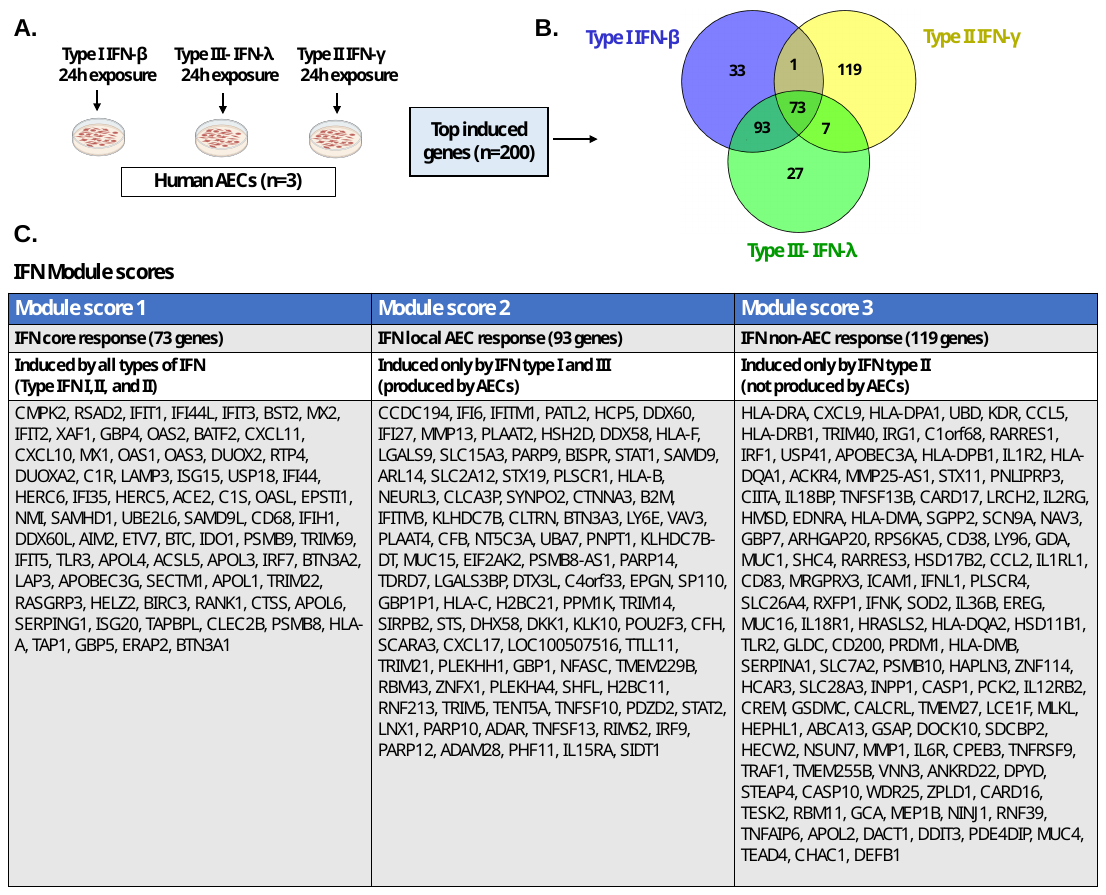
**Supplementary Figure. 1. Generation of IFN module scores. A**. Nasal AEC from human infant donors (n=3) were exposed to type I IFN/β, type II IFN/γ or type III IFN/λ and then transcriptomic profiles (RNA-seq) were obtained. **B.** Venn diagram showing overlap of the top 200 genes of each condition. **C.** Module scores and corresponding genes included in each condition.

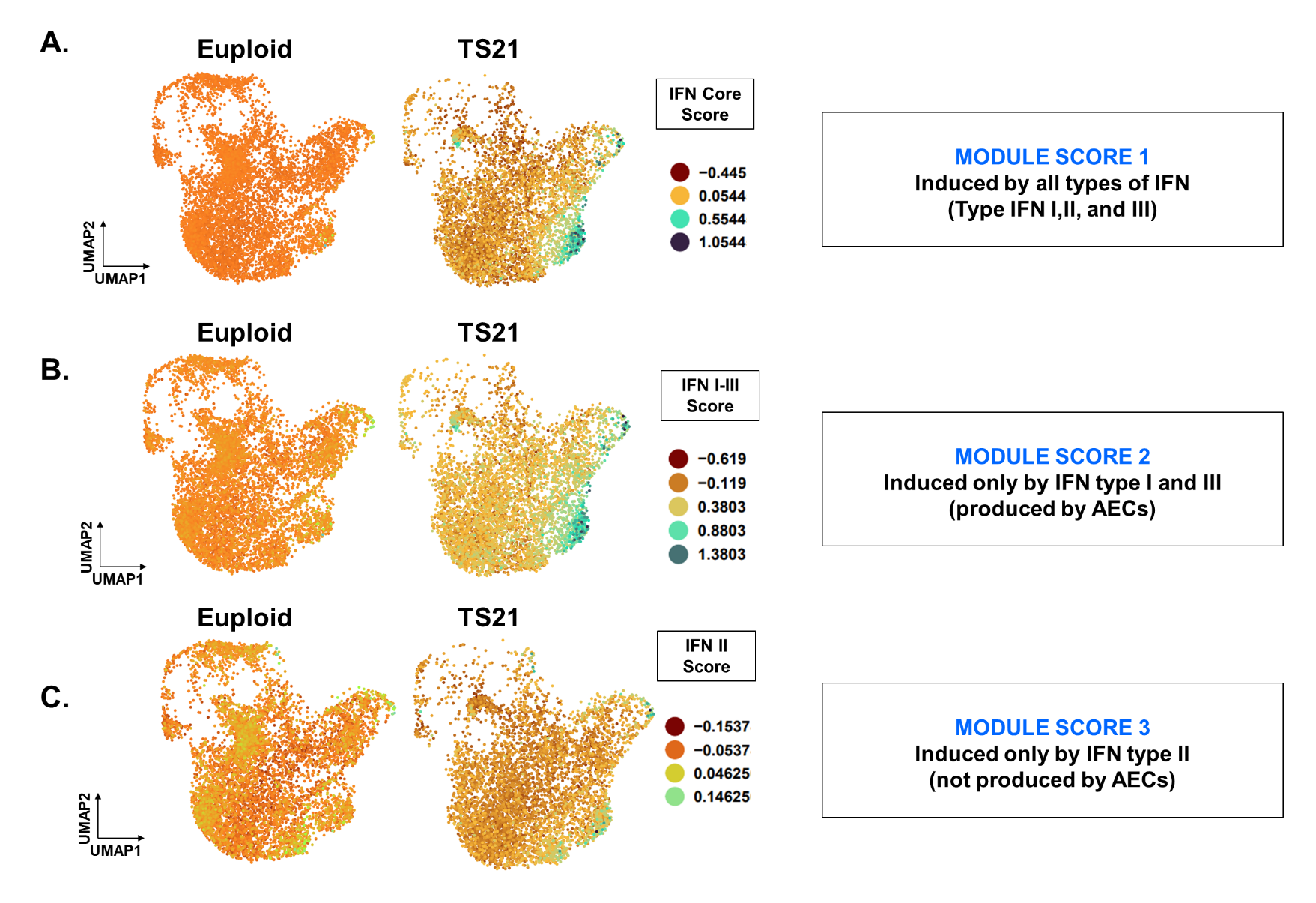

**Supplementary Figure. 2. IFN module scores in airway epithelial cells. A-C.** UMAP of scRNA-seq data shows a subpopulation of TS21 AECs that have upregulation of IFN-inducible genes (marked in green) identified using a module score encompassing core IFN genes induced by all type of IFN **(A)** or those induced by type I-III IFN **(B)** but not by a module score induced by type II IFN only **(C).** Results representative of uninfected AECs from an infant donor with DS (n=5,742 cells) and an age-matched euploid donor (n=7,879 cells).

**Supplementary Figure 3. Type-I IFN response is suppressed in human airway epithelial cells.** Type-I IFN/β protein levels at baseline (labelled as “0”) and during RSV infection (MOI 1 and MOI 3 x24 hours). Results representative of AECs derived from n=6 euploid donors (EUP) and n=6 individuals with DS (TS21).
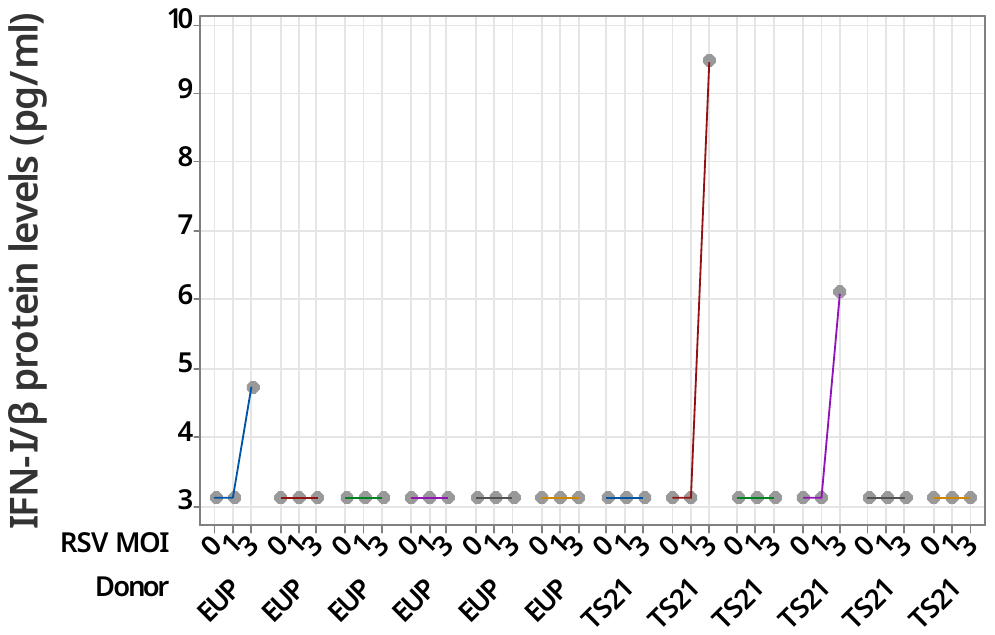
Values were below level of detection (3.1 pg/ml) in most donors.
